# Large-Scale COI Gene Data Analysis for the Development of a Gel-Based Multiplex PCR Test Capable of Identifying Biting Midge Vector Species and Haplotypes (Diptera: Ceratopogonidae) of the *Culicoides* subgenus *Avaritia* Fox, 1955

**DOI:** 10.1101/2024.01.23.576915

**Authors:** Oliver Dähn, Doreen Werner, Bruno Mathieu, Helge Kampen

**Affiliations:** Friedrich-Loeffler-Institut (FLI), Federal Research Institute for Animal Health, Südufer 10, Greifswald – Insel Riems 17493, Germany; Leibniz Centre for Agricultural Landscape Research (ZALF), Eberswalder Str. 84, Müncheberg 15374, Germany; Institutes of Bacteriology and Parasitology of the Medical Faculty, University of Strasbourg, UR 3073 PHAVI, 67000 Strasbourg, France

**Keywords:** *Culicoides*, Obsoletus Group, *Avaritia*, vector, polymerase chain reaction (PCR), mitochondrial cytochrome c oxidase subunit I (COI), haplotype

## Abstract

The emergence of culicoid-transmitted bluetongue and Schmallenberg viruses in several European countries demonstrated the ability of indigenous biting midge species to transmit pathogens. Entomologic research programmes identified members of the Obsoletus Group (*Culicoides* subgenus *Avaritia*) as keyplayers in disease epidemiology in Europe. However, morphological identification of potential vectors to species level is challenging due to the existence of isomorphic species. PCR tests developed to facilitate genetic species determination have been dismantled by the discovery of new genetic variants (haplotypes) of *C. obsoletus* sensu stricto (s.s.), forming distinct clades. In this study, 4,422 GenBank entries of the mitochondrial cytochrome c oxidase subunit I (COI) gene of subgenus *Avaritia* members of the genus *Culicoides* were analyzed to develop a conventional multiplex PCR, capable of detecting all vector species and recently described clades of the western Palaearctic in this subgenus. Numerous GenBank entries incorrectly assigned to a species were identified, analyzed and reassigned. The results suggest that the three clades of *C. obsoletus* represent independent species, whereas *C. montanus* should rather be regarded as a genetic variant of *C. obsoletus* s.s.. Based on these findings, specific primers were designed and validated with DNA material from field-caught biting midges which achieved very high diagnostic sensitivity (100%) when compared to an established reference PCR (82.6%). Hence, the newly developed multiplex PCR represents the first molecular tool which enables both the identification of accepted species and of the three clades of *C. obsoletus* s.s. and could provide new insights into the ecology of the latter.

## Introduction

With a size of merely 1-3 mm, culicoid biting midges (Diptera: Ceratopogonidae) belong to the smallest haematophagous insects on the planet, colonizing almost all continents and climate zones, except Antarctica and New Zealand [1]. More than 50 different arboviruses and a variety of parasitic nematodes and protozoans are known to be transmitted by *Culicoides* species. With their widespread distribution and high population sizes, biting midges are important vectors of disease agents of ruminants, equids and – in rare cases – humans [1,2,3].

In Europe, bluetongue virus (BTV) and African horse sickness virus (AHSV) were isolated from specimens of the Obsoletus Group (subgenus *Avaritia* Fox, 1955) and the Pulicaris Group (subgenus *Culicoides* Latreille, 1809) more than 40 years ago [4,5]. Notwithstanding, little scientific attention has been given to indigenous biting midge species as vectors of disease agents until the unprecedented outbreak of bluetongue (BT) in Central Europe in 2006. Surprisingly, this occurred in areas where the classical Afro-Asian vector species of BTV, *C. imicola* Kieffer, 1913, has never been recorded. Thus, the BT epidemic suggested vector competence of European biting midges, and subsequent entomological studies confirmed long-suspected involvement of species of the Obsoletus and Pulicaris Groups in disease epidemiology [6–14]. A few years later, species from these groups were also linked to the newly emerged Schmallenberg virus (SBV) [15–26], and it is conceivable that they could be competent vectors of other viruses as well. So far, evidence of vector competence of endemic species is scarce, but preliminary infection studies [27,28] and frequent virus isolation from field-caught Obsoletus Group biting midges appear to confirm their involvement in the transmission of both BTV and SBV.

Adult *Culicoides* are commonly identified on the basis of phenotypic characteristics, such as wing pigmentation, following published identification keys [29–32]. However, this approach is time consuming, requires many years of experience and is inefficient in analyzing large quantities of midges [33]. The discovery of closely related species with similar or identical morphology further complicated traditional species identification and made it necessary to implement taxonomic categories below the subgenus level, such as ’Obsoletus Group’ [2,34,35]. Nine valid species are currently assigned to this group, namely *C. abchazicus* Dzhafarov, 1964; *C. alachua* Jamnback and Wirth, 1963; *C. filicinus* Gornostaeva and Gachegova, 1972; *C. gornostaevae* Mirzaeva, 1984; *C. montanus* Shakirzjanova, 1962; *C. obsoletus* (Meigen, 1818); *C. sanguisuga* (Coquillett, 1901); *C. scoticus* Downes and Kettle, 1952; and *C. sinanoensis* Tokunaga, 1937 [36]. For additional subclassification within the Obsoletus Group, Meiswinkel et al. [37] introduced the term ’Obsoletus Complex’ or ’Obsoletus/Scoticus Complex’, which groups closely related species, whose isomorphic females cannot be distinguished morphologically. This complex includes *C. obsoletus*, *C. montanus* and *C. scoticus*.

The confusion about biting midge taxonomy is further increased by the commonly accepted use of synonyms for one and the same species as a result of simultaneous description of specimens in different regions and multiple denominations. For instance, the most frequent and eponymous species of the Obsoletus Group, *C. obsoletus*, had also been described as *C. varius* (Winnertz, 1852); *C. yezoensis* (Matsumura, 1911); *C. lacteinervis* Kieffer, 1919; *C. rivicola* Kieffer, 1921; *C. clavatus* Kieffer, 1921; *C. heterocerus* Kieffer, 1921; *C. pegobius* Kieffer, 1922; *C. kabyliensis* Kieffer, 1922; *C. concitus* Kieffer, 1922; *C. intermedius* Okada, 1941; *C. sintrensis* Cambournac, 1956; and *C. seimi* Shevchenko, 1967 [38]. However, by and by all these names have been officially synonymised under the species name ’*C. obsoletus*’, according to the guidelines of the International Code of Zoological Nomenclature (ICZN).

Based on the mentioned issues, morphological species determination has increasingly been replaced or supplemented by molecular methods such as DNA barcoding. These provided a deeper insight into the genetics of biting midges but, at the same time, raised concern about assumed phylogenetic relationships, particularly for isomorphic taxa. For instance, two further species of the West Palaearctic fauna, *C. chiopterus* (Meigen, 1830) and *C. dewulfi* Goetghebuer, 1936, had been considered part of the Obsoletus Group/Obsoletus Complex for many years [39]. However, already in 2008, a study investigating DNA sequence variation of the mitochondrial cytochrome c oxidase subunit I (COI) gene and the ribosomal internal transcribed spacers (ITS1 and ITS2) postulated that *C. dewulfi* be separated from the Obsoletus Complex [40], which was later supported by other authors [41–44]. Another study went one step further and combined molecular analyses with morphometric measurements and unequivocally excluded *C. chiopterus* and *C. dewulfi* from the Obsoletus Group [35], whereas the results of a later genetic study on *Culicoides* relationships, using Bayesian inference analysis, re-located *C. chiopterus* in the group [43]. Additionally, further molecular work demonstrating the genetic distance between *C. montanus* and *C. obsoletus* to be similar to intraspecific distances between different *C. obsoletus* specimens questioned whether *C. montanus* should be regarded a true separate species of the group or merely a genetic variant of *C. obsoletus* [36,45].

The discovery of previously unknown genetic variants of *C. obsoletus* from Sweden and Switzerland, denoted as *C. obsoletus* clades O1, O2 and O3 [41,46], as well as *C. obsoletus* ’clade dark’ from the Netherlands [37,47] and two phylogenetic clades (1 and 2) of *C. scoticus* from several European countries [36,43] significantly increased the number of Obsoletus Group members and clearly illustrated the taxonomic complexity of the subgenus. A comprehensive multi-marker study trying to arrange these genetic variants within the Obsoletus/Scoticus Complex, concluded that sequences named *C. obsoletus* clade O1 actually represent *C. obsoletus* sensu stricto (s.s.) and *C. obsoletus* ‘clade dark’ is a synonym of *C. obsoletus* clade O3 [36]. Another group of authors hypothetized that *C. obsoletus* ‘clade dark’ could be *C. gornostaevae* [37], a boreal species which was first described only from Siberia, but has recently been reported from Norway, Poland and Sweden [48].

In previous studies, the COI gene proved to be a suitable marker for the differentiation of species of several insect groups, including mosquitoes [49,50], sandflies [51], gallwasps [52] and tabanids [53]. Subsequently, COI barcoding has successfully been used for species determination of culicoid biting midges as well, particularly within the *Culicoides* subgenera *Culicoides* [41,54] and *Avaritia* [41,55–57], as it is a sufficiently long, high copy gene that is composed of both conserved and variable regions [58–60].

However, this method cannot be applied on pooled samples due to the risk of mixed or impure sequences. Especially, if multiple species are present in a pool, the generated sequence is likely to belong to the most abundant one or, if one or more midges are freshly engorged, DNA barcoding might rather determine the blood-host species [61] due to the high volume of ingested blood relative to the small size of the insect [62].

Several PCR tests have been developed to address identification problems, capable of differentiating females of closely related species within the *Culicoides* subgenus *Avaritia* with very similar or identical morphologic features and juvenile specimens (larvae, pupae) in which those distinguishing features are not yet developed [46,63–70]. However, these PCR tests were developed many years ago, often using a small and geographically restricted gene pool, which significantly limits the applicability of the tests. Additionally, the genetic variants of *C. obsoletus* and *C. scoticus* were not considered for PCR development and are therefore not detectable with those tests.

Hence, an intensive GenBank analysis was carried out in the present study in order to (i) clarify the taxonomic status of the different valid and cryptic species within the *Culicoides* subgenus *Avaritia* and use these findings to (ii) develop an easy-to-use multiplex PCR assay for the identification of putative BTV and SBV vector species of the West Palaearctic biting midge fauna, including *C. scoticus* (clades 1 and 2), *C. dewulfi*, *C. chiopterus* and the recently described genetic variants of *C. obsoletus* (clades O1, O2 and O3) whose habitat preferences and roles in disease transmission are yet unknown.

## Materials and Methods

### Insect Collection and Morphologic Examination

The majority of culicoid biting midges were collected with BG-Sentinel UV-light suction traps (Biogents, Regensburg, Germany) during various monitoring activities in Germany. Few individuals originated from samplings in other European countries, Russia and USA. Captured midges were morphologically pre-identified to group or species level using a stereomicroscope and common identification keys [29–32] and kept in 75% EtOH for molecular analysis.

### DNA Extraction

Single specimens of selected insects were removed from the storage vessel and put onto a clean paper tissue. After evaporation of fixative residues for 1 min at room temperature, they were transferred to Eppendorf tubes containing either 180 µL buffer ATL and 20 µL Proteinase K (Qiagen, Hilden, Germany), or 350 µL of in-house ZB5d medium/antibiotics mixture as described in Ries et al. [71]. Insects were homogenized for 3 min at 30 Hz with TissueLyser II (Qiagen), using three 3 mm steel beads (TIS GmbH, Gauting, Germany). Total genomic DNA was subsequently isolated from the ATL/Proteinase K mixture with the QIAamp DNA Mini Kit (Qiagen, Hilden, Germany), according to the manufacturer’s instructions, in 50 µL elution volume, while the ZB5d/antibiotics mixture was further processed with the NucleoMag VET Kit (Macherey-Nagel, Düren, Germany) in a final elution volume of 100µL VEL-buffer using a KingFisher®Flex automat (Thermo Fisher Scientific, Dreieich, Germany).

### COI Amplification and Sequencing

Isolated DNA was used to generate partial fragments of the mitochondrial (mt) COI gene either with species-specific primers or universal primers PanCuli-COX1-211F and PanCuli-COX1-727R as described in Lehmann et al. [70]. Additionally, a newly designed generic primer PanCuli-COX1-025F was used instead of PanCuli-COX1-211F, to amplify a longer fragment of the COI gene according to the protocol of Dähn et al. [61]. Produced PCR products were mixed with 2.5 µL of 6x DNA loading dye (Thermo Fisher Scientific) and analyzed on 1.5% agarose gels, which had been supplemented with 5 mg/mL of ethidium bromide solution and were run for 50 min at 100 V. The gels were visualized with a ChemiDoc MP Imaging System (Bio-Rad, Feldkirchen, Germany), and amplicons of expected lengths were excised and extracted with the QIAquick Gel Extraction Kit (Qiagen). For sequencing, DNA fragments were cycle-sequenced with one of the PCR primers using the BigDye Termintor v1.1 Cycle Sequencing Kit (Thermo Fisher Scientific). Resulting PCR products were purified with the Bioanalysis NucleoSEQ Kit (Macherey-Nagel), and 15 µL of each eluate was mixed with the same volume of Hi-Di formamide (Thermo Fisher Scientific). Samples were finally sequenced either in one or in both directions on a 3500 Genetic Analyzer device (Applied Biosystems/Hitachi, Darmstadt, Germany). Obtained sequences were edited with Geneious Prime software version 2021.0.1 (Biomatters, Auckland, New Zealand) and checked against NCBI GenBank (www.ncbi.nlm.nih.gov). Edited sequences were deposited in GenBank and corresponding DNA samples were later used for PCR validation.

### Data Analysis and Primer Design

Initially, the GenBank was browsed for all available COI, ITS1 and ITS2 sequences of the different Obsoletus Group members (valid species and genetic variants) including *C. chioperus* and *C. dewulfi* and their synonyms found in the literature. Since different designations of the genes existed, alternative terms were also considered. However, due to the lack of genetic data for the other gene loci, only the COI gene provided a sufficient number of sequences of the respective species and haplotypes to design specific primers.

Collected COI sequences were checked for plausibility through comparison of all published GenBank entries available for the Obsoletus Group taxa in a Geneious multiple alignment. The determined genetic distances between individual sequences were further used for Microsoft Excel (Microsoft Corporation, USA) calculations to identify incorrect entries (**Supplementary Table S1**). Assuming that the majority of GenBank entries (more than 50%) were accurately assigned and sequences at least 98.5% similar to each other belong to the same species or haplotype, a factor for each GenBank entry (here named ‘species determination factor’ or SDF) was calculated according to the following equation:

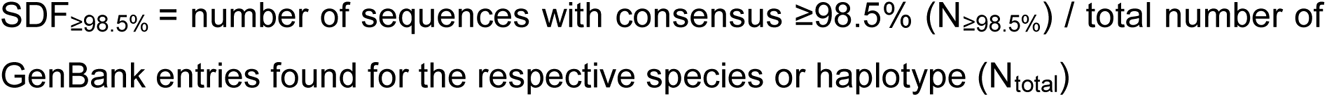

Following this formula all entities with SDF_≥98.5%_ > 0.5 were considered to belong to the respective species or haplotype. In contrast, sequences with SDF_≥98.5%_ < 0.5 were categorized as ‘dubious’, re-analyzed with NCBI nucleotide BLAST tool (https://blast.ncbi.nlm.nih.gov/Blast.cgi, last data request on 15 November 2022) and assigned to matching taxa or – if no sequence match could be found – excluded from further analysis. Selected sequences (**Supplementary Table S1**) were used to create species- or haplotype-specific consensus sequences with Geneious Prime software, which were finally compared in a Geneious multiple alignment using initial settings. Inter- and intraspecific variances in DNA sequence were used to design specific forward primers, according to common guidelines for primer design [72–74]. Selected primer candidates were checked regarding melting temperature, GC-content, self-dimerization and primer-dimer formation using Oligo Analysis Tool (https://eurofinsgenomics.eu/en/ecom/tools/oligo-analysis/, last access on 11 December 2022) and analyzed with NCBI nucleotide BLAST tool for repetitive sequences. Primers were checked for functionality, specificity and the capability of multiplexing with molecularly pre-identified biting midge DNA or—in case no DNA material of the respective taxon was available or a defined amount of DNA had to be used—synthetic COI gene DNA (**Supplementary Table S2**) produced by GenExpress (Berlin, Germany).

### Performance of Multiplex PCR

The newly designed forward primers were applied in a single-tube multiplex PCR in combination with the universal reverse primer PanCuli-COX1-727R [70]. The master mix was composed of 10 µL 2x QuantiTect Multiplex PCR NoROX reagent (Qiagen), 0.5 µM of each primer and 2 µL DNA template, and supplemented with ddH_2_O to give a total volume of 20 µL.

In order to enable simultaneous examination of morphologically pre-identified specimens of the subgenera *Avaritia* Fox and *Culicoides* Latreille, DNA amplification was performed using the same cycling conditions as mentioned in Dähn et al. [61]: 15 min at 95 °C (activation of Taq polymerase), followed by 42 cycles of 30 s at 95 °C (denaturation), 45 s at 63 °C (primer annealing), 45 s at 72 °C (primer extension), and a final elongation step for 5 min at 72 °C. The PCR products were subsequently applied to agarose gels and visualized as described. The developed multiplex PCR was validated with genetically pre-identified (sequenced) DNA of field-collected biting midges and several non-ceratopogonid insect species, and results were compared to those obtained by a previously published reference PCR test [70].

## Results

### GenBank Search and Data Analysis

In the present study, searching GenBank initially proved to be very difficult due to the use of synonyms for one and the same species and different nominations of the gene, e.g. ‘COX1’, ‘COI’, ‘CO1’ or the unabbreviated term ‘mitochondrial cytochrome c oxidase subunit I’.

With 4,422 sequences found, the COI gene was by far the most common marker gene deposited in GenBank. Except for *C. gornostaevae* and *C. filicinus*, for which no COI sequences were available at the time of browsing (last data request: 15 November 2022), sequences were collected for all species and haplotypes and checked for plausibility (**Supplementary Table S1**). The majority of the published sequences (n=2,796, 63.2%) were assigned to *C. obsoletus*, with 91.7% of them showing ≥98.5% similarity to each other in more than 50% of all GenBank entries designated as ‘*C. obsoletus*’. The remaining 233 COI sequences (8.3%) were dubiously different from the other sequences and could be assigned to *C. scoticus* clade 1 (30.9%), *C. sanguisuga* (23.2%), *C. obsoletus* clade O2 (19.3%), *C. obsoletus* clade O3 (8.6%), *C. montanus* (6.4%) and *C. scoticus* clade 2 (0.9%). For 25 of the dubious sequences (10.7%), no distinct assignment could be made with the used calculation model, however, 68% of these entities showed highest similarity to *C. sanguisuga* (96.5-98.7%) and 32% showed highest similarity to *C. obsoletus* clade O1 (96.8-100%).

For *C. obsoletus* clade O2, 519 COI sequences were found in GenBank, with 10.0% of them being identified as incorrectly allocated to this haplotype. Those could be reassigned to *C. obsoletus* clade O1 (78.8%), *C. scoticus* clade 1 (11.5%), *C. scoticus* clade 2 (5.8%) and *C. obsoletus* clade O3 (3.8%).

In the case of *C. obsoletus* clade O3, only six COI sequences could be found in GenBank with this designation. However, the re-analysis of dubious sequences of other subgenus *Avaritia* members and browsing for the synonymous name ‘*Culicoides* sp. ’dark obsoletus’ sensu Meiswinkel et al. (2015)’ [37] identified 47 additional sequences with high pairwise identity (98.7-100%) to the sequences designated as *C. obsoletus* clade O3.

The second highest number of all analyzed subgenus *Avaritia* species and haplotype sequences were found for *C. scoticus* clade 1 (811 of 4,422 sequences). Critical cross-checking of these sequences identified 19.5% as incorrectly assigned, actually belonging to *C. scoticus* clade 2 (n=75, 47.5%), *C. obsoletus* clade O1 (n=74, 46.8%), and *C. obsoletus* clade O2 (n=9, 5.7%). Additionally, two ‘*C. scoticus*’ sequences (MT172703 and MT172804) showed highest pairwise identity to *C. scoticus* clade 1 (96.6-98.4), however, they did not reach the 98.5% consensus value and were therefore excluded from further analysis. Sequences of the latter haplotype, *C. scoticus* clade 2, could not be found in GenBank under the respective notation. However, the comparison of dubious sequences identified in the framework of this study with sequences of specimens designated as ‘*C. scoticus* clade 2’ from Mignotte et al. [36] revealed a considerable number of COI sequences of this genetic variant (n=74) collected in nine different countries. Moreover, 116 COI sequences of *C. montanus*, a taxa which has already been discussed by some authors to be a genetic variant of *C. obsoletus*, were found in GenBank, with most of them (n=79, 68.1%) originating from specimens caught in Morocco. Three of the 116 GenBank entries (2.6%) designated as ‘*C. montanus*’ were found to be incorrectly deposited and could be assigned to other Obsoletus Complex members (*C. obsoletus* clade O1: n=1, *C. scoticus* clade 1: n=2).

For the remaining *Culicoides* subgenus *Avaritia* members, only ten COI sequences were found in total: *C. sinanoensis* (n=5), *C. sanguisuga* (n=2), *C. abchazicus* (n=2), *C. alachua* (n=1); but no sequences were available for *C. gornostaevae* and *C. filicinus*. The low number of sequences found for these species made it extremely difficult to verify their integrity, and it had to be trusted that these specimens were identified correctly. Considering this, 53 COI sequences from *Culicoides* collected in Canada (deposited as ‘*C. obsoletus*’), showed very high similarity to the sequences of *C. sanguisuga* (MK760237 and MK760238) and could therefore be successfully assigned to this species. In case of the five sequences of *C. sinanoensis*, the data analysis identified two sequences as ‘dubious’, but the re-analysis suggested that they might be genetic variants of this species due to comparatively high consensus (98.1-98.4%) to the sequences confirmed as *C. sinanoensis*.

In case of the two species once included in the Obsoletus Group, *C. chiopterus* showed comparatively low intraspecific pairwise identity (12 of 80 sequences were significantly different and were therefore excluded), whereas all sequences representing ‘*C. dewulfi*’ (n=84) were quite similar to each other (98.7-100%) and seemed to be correctly designated. However, all of the dubious sequences of *C. chiopterus* were closely related to the sequences confirmed to belong to this species and five of them (HQ824396, HQ824402, HQ824403, JQ898002, JQ898006) seem to form a monophyletic clade (pairwise identity: 99.8%) which is 1.2% divergent from the consensus sequence of *C. chiopterus*.

In summary, of the 4,422 COI sequences found in GenBank, 4,407 could accurately be designated to species or haplotype (**Figure 1**). With exception of the two *C. abchazicus* sequences, which were only 97.4% similar to each other, generated consensus sequences showed very high intraspecific (pairwise) identities of 99.4% to 99.9%, indicating the accuracy of sequence reassignment. However, multiple alignment comparison of generated consensus sequences revealed very low interspecific divergence in COI sequence for *C. obsoletus* clade O1 and *C. montanus* (2.8%), which was comparable to that of the recently discovered variants (clade 1 and clade 2) of *C. scoticus* (2.9%), questioning its taxonomic status as separate species. By contrast, the interspecific divergence of the three clades of *C. obsoletus* was comparatively high (O1 vs. O2: 10.4%, O1 vs. O3: 10.1%, O2 vs. O3: 9.3%). Additionally, the high interspecific divergence of *C. dewulfi* from the members of the Obsoletus Group (16.6% to 21.2%, average: 18.9%) seem to confirm its distinct status outside the group, whereas the mean interspecific divergence of *C. chiopterus* (15.7%) from the valid species and haplotypes of the Obsoletus Group, was exactly between *C. alachua* (12.6%), an accepted member of the group, and *C. dewulfi*.

**Figure 1.**
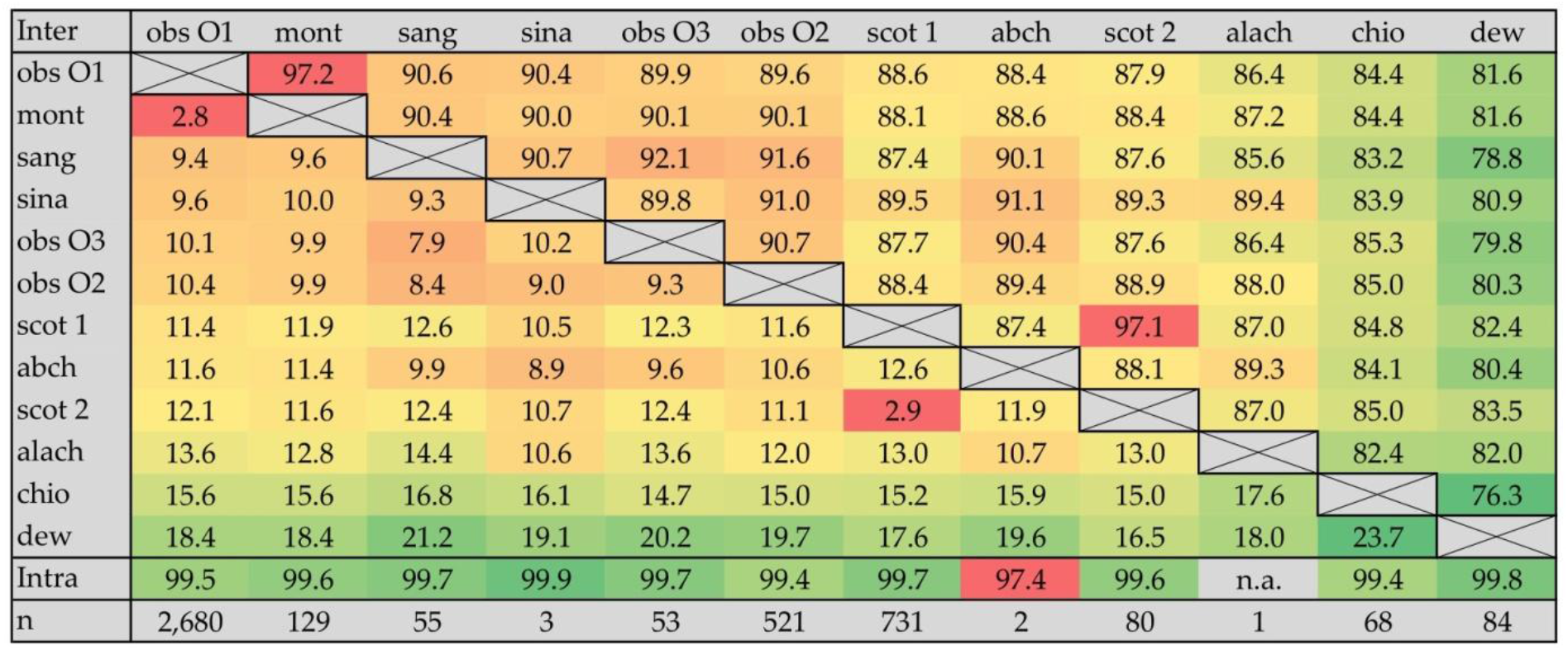
Inter- and intraspecific pairwise comparison of COI gene DNA sequences between the analyzed taxa of the *Culicoides* subgenus *Avaritia*: Interspecific genetic distances are displayed in the left-bottom half of the matrix and highlighted with graded colors from red (low distance) through yellow (medium distance) to green (high distance). Interspecific pairwise identities in gene sequence are presented in graded colors in the right-upper half of the matrix with the opposite meaning of the colors: red (high similarity) – yellow (medium similarity) – green (low similarity). Intraspecific pairwise identities (Intra) are given as well, using the same color code. Values (in %) were calculated through the comparison of species- and haplotype-specific consensus sequences of respective GenBank entries (n). *C. obsoletus* clade O1 (obs O1), *C. montanus* (mont), *C. sinanoensis* (sina), *C. obsoletus* clade O3 (obs O3), *C. obsoletus* clade O2 (obs O2), *C. sanguisuga* (sang), *C. scoticus* clade 1 (scot 1), *C. abchazicus* (abch), *C. scoticus* clade 2 (scot 2), *C. alachua* (alach), *C. chiopterus* (chio) and *C. dewulfi* (dew).

### Primer Design and PCR Performance

Multiple alignment comparison of generated consensus sequences revealed genetic differences, which were used to design specific forward primers for selected taxa of the subgenus *Avaritia* according to the PCR concept of Lehmann et al. [70]. The focus was on putative vector species of the West Palaearctic biting midge fauna, including *C. obsoletus* clade O1, *C. obsoletus* clade O2, *C. obsoletus* clade O3, *C. scoticus* (clades 1 and 2), *C. chiopterus* and *C. dewulfi*. Since low interspecific COI gene sequence divergence of *C. montanus* from *C. obsoletus* clade O1 (2.8%) and of *C. scoticus* clade 1 from *C. scoticus* clade 2 (2.9%), respectively, makes the development of specific primers almost impossible, no specific primers could be designed for these four taxa.

Of 52 forward primers tested (**Supplementary Table S3**), many revealed cross-reactivity to other members of the Obsoletus Group or unsuitability for multiplexing. However, length variation and integration of wobble-bases improved both parameters, except in case of the forward primer designated for *C. dewulfi*, which needed additional, site-directed insertion of a mismatch base to guarantee primer specificity. The best-working forward primers (**Table 1**) were pre-tested regarding their functionality (detection of the specific target DNA) and specificity (reactivity with DNA of the other *Culicoides* taxa considered for PCR development) (**Figure 2A-F**), as well as their suitability for multiplexing (**Figure 2G**).

**Figure 2.**
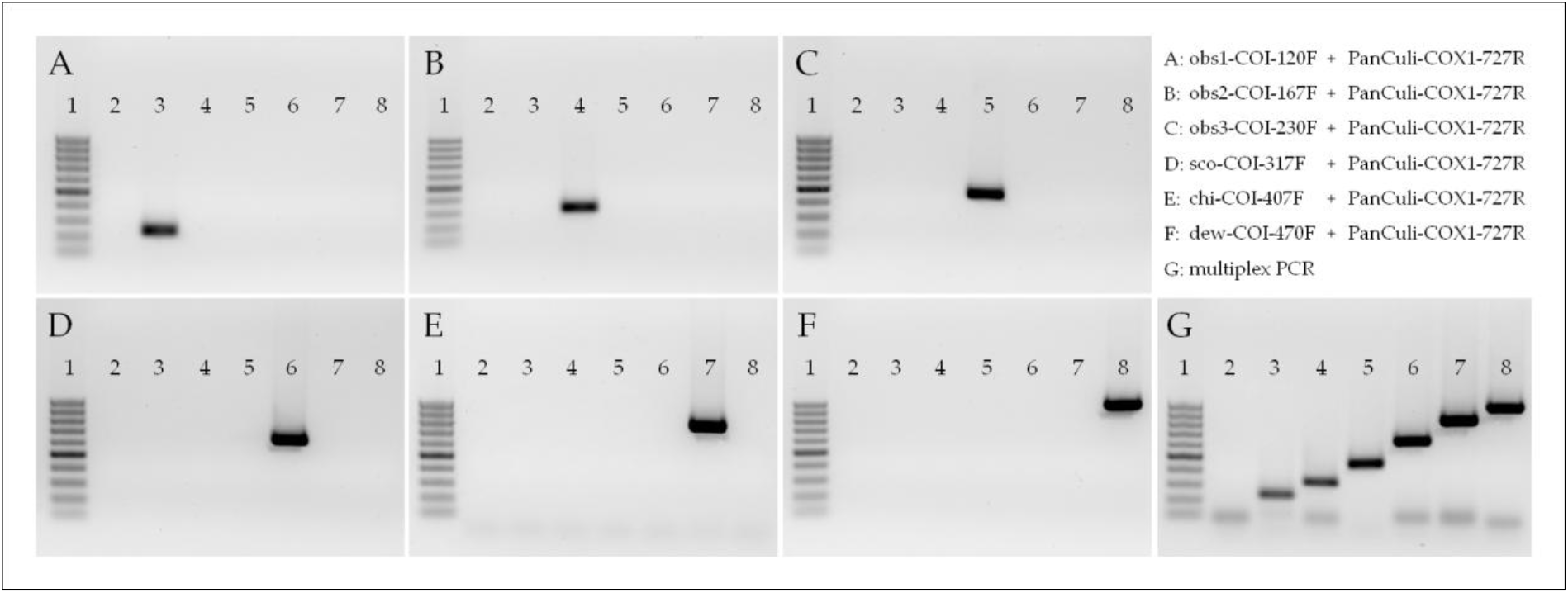
Proof of function of the multiplex PCR test for the members of the Obsoletus Group, including *C. chiopterus* and *C. dewulfi*. Specific forward primers were tested regarding their specificity (singleplex, A-F) and capability for multiplexing (G). The forward primers used were the following: obs1-COI-120F (A, G), obs2-COI-167F (B, G), obs3-COI-230F (C, G), sco-COI-317F (D, G), chi-COI-407F (E, G) and dew-COI-470F (F, G). All primers were used in combination with the universal reverse primer PanCuli-COX1-727R. DNA samples used for PCR validation contained either 106 copies of specific target or 107.5 copies of unspecific target (synthetic COI gene). For *C. dewulfi* and *C. chiopterus*, equivalent amounts of quantified COI gene amplicon were used. Lane 1: 50 bp ladder (50-500 bp Gene Ruler; Roth, Karlsruhe, Germany), lane 2: no template control, lane 3: *C. obsoletus* clade O1, lane 4: *C. obsoletus* clade O2, lane 5: *C. obsoletus* clade O3, lane 6: *C. scoticus* clade 1, lane 7: *C. chiopterus* and lane 8: *C. dewulfi*.

**Table 1.**
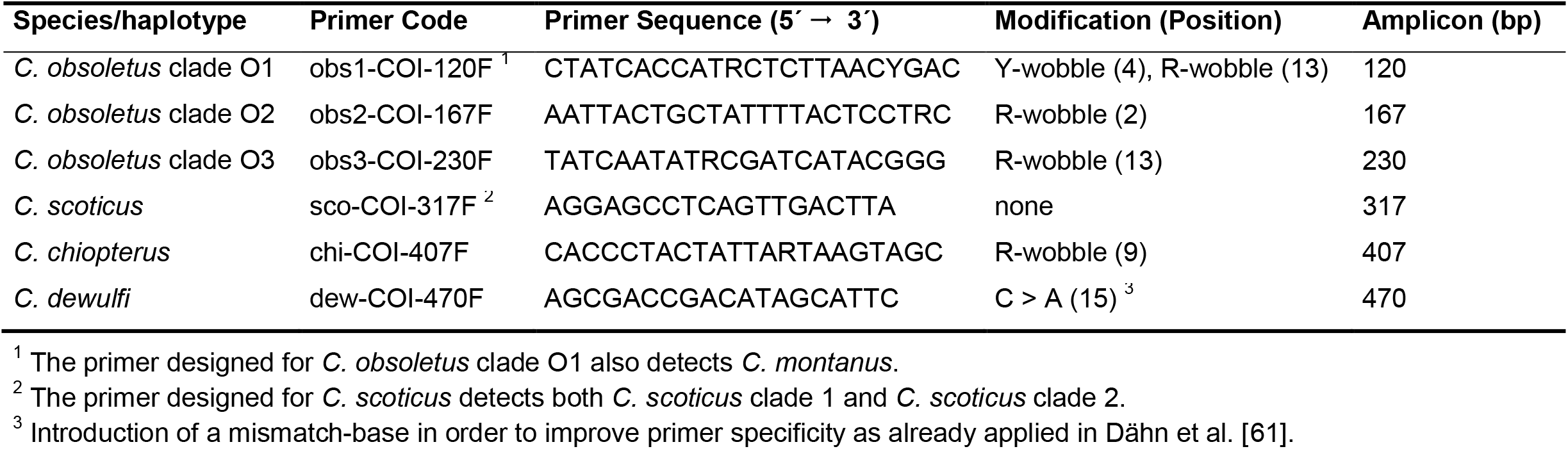
List of specific forward primers designed for the four members of the Obsoletus Group plus *C. dewulfi* and *C. chiopterus*. The primers can be combined in a single-tube multiplex approach using the universal primer PanCuli-COX1-727R as a reverse primer.

To confirm that even one single biting midge (equivalent to approximately 10^6^ copies of mt genome) can reliably and specifically be detected in a pool of 50 individuals, 50 times higher amounts (10^7.5^ copies) of synthetic COI gene of the unspecific target as compared to the specific DNA were used. In this test, the specific forward primers, in combination with the universal reverse primer PanCuli-COX1-727R, showed the expected amplicon lengths between 120 bp and 470 bp (**Figure 2**) for single specimens of the target species (10^6^ copies of synthetic DNA), without detecting 10^7.5^ copies of unspecific DNA. Furthermore, no amplification was observed in no-template negative controls (**Figure 2, lane 2**).

### Validation of the Multiplex PCR

After successful pre-testing, the developed multiplex PCR was validated regarding sensitivity and specificity by testing of genomic DNA from field-caught biting midges (or synthetic DNA for specimens not found in field-collections) and direct comparison of the results to those achieved with the same DNAs in a reference PCR test. To ensure the precision of the results and eliminate uncertainties, the DNA test samples underwent prior verification by COI barcoding.

When evaluating the primers for their diagnostic sensitivity against genomic DNA of 92 single biting midge specimens, the newly developed multiplex PCR (mPCR) detected all specific DNAs, giving a total diagnostic sensitivity of 100% (**Table 2**). By contrast, the published reference PCR [70] exhibited notable limitations, particularly concerning the identification of *C. obsoletus* clade O2 (53.6%), *C. obsoletus* clade O3 (66.7%) and *C. scoticus* (80.0%), gaining a total sensitivity of only 82.6%.

**Table 2.** Determination of diagnostic sensitivity of the developed mPCR. A total of 92 samples belonging to the Obsoletus Group (plus *C. dewulfi* and *C. chiopterus*) were tested with the newly developed multiplex PCR and compared to the results achieved with a reference PCR [70]. DNA extracts of genetically pre-identified, single specimens were used for testing.

| Species/haplotype | GenBank accession no. | Specimens tested [n] | New mPCR |  | Reference mPCR <sup>1</sup> |  |
| --- | --- | --- | --- | --- | --- | --- |
|  |  |  | Positive<br>[n] | Sensitivity<br>[%] | Positive<br>[n] | Sensitivity<br>[%] |
| *C. obsoletus* clade O1 | OQ789075, OQ941500-537 | 39 | 39 | 100 | 39 | 100 |
| *C. obsoletus* clade O2 | OQ789076, OQ941538-560, PP110209-212 | 28 | 28 | 100 | 15 | 53.6 |
| *C. obsoletus* clade O3 | OQ789077, OQ941561-562 | 3 | 3 | 100 | 2 | 66.7 |
| *C. scoticus* clade 1 | OQ941563-572 | 10 | 10 | 100 | 8 | 80.0 |
| *C. chiopterus* | KJ624070, MK760108, MK760110, OQ789068, OQ941573 | 5 | 5 | 100 | 5 | 100 |
| *C. dewulfi* | MK760112-114, OQ789069, OQ941574-576 | 7 | 7 | 100 | 7 | 100 |
| Total |  | 92 | 92 | 100 | 76 | 82.6 |

<sup>1</sup> The reference test is not able to distinguish between the different clades of *C. obsoletus* since a single primer for *C. obsoletus* (obs-COI-fwd) is used.

In the course of specificity determination, 36 other *Culicoides* species and haplotypes were tested, belonging to seven different subgenera, namely *Avaritia* Fox (n=5), *Beltranmyia* Vargas (n=1), *Culicoides* Latreille (n=20), *Monoculicoides* Khalaf (n=1), *Sensiculicoides* Shevchenko (n=6), *Silvaticulicoides* Glukhova (n=1), *Wirthomyia* Vargas (n=1) and one species (*C. pallidicornis*) unplaced into a subgenus (**Table 3**). In addition to the subgenus *Avaritia*, the focus of specificity testing was on members of the *Culicoides* subgenus *Culicoides*, since those are also frequently found in field catches of the West Palaearctic region. Due to difficulties in acquisition of respective DNA material within this subgenus, synthetic COI gene DNA was used in some cases. The primers of both the newly developed PCR and the reference PCR revealed cross-reactivity to several subgenus *Culicoides* species and haplotypes. In the case of the newly developed mPCR, unspecific amplification of the non-target species was observed for 15 out of 36 samples, whereby most false-positive results (n=9) were achieved with the *C. scoticus* forward primer (sco-COI-317F), mainly with DNA of the *Culicoides* subgenus *Culicoides* (n=6). As already assumed due to few COI sequence differences, the genomic DNA of *C. scoticus* clade 2 was incorrectly detected with the primer sco-COI-317F and the *C. montanus* sample with the primer of *C. obsoletus* clade O1 (obs1-COI-120F). The primers of the reference PCR exhibited comparable deficiencies, leading to the inaccurate detection of 11 out of 36 ‘other *Culicoides*’ (taxa not belonging to the subgenus *Avaritia*) and in certain instances, samples were even falsely identified with two non-specific primers (**Table 3**). Most cross-reactivity was observed with the *C. obsoletus* (obs-COI-fwd)-specific primer, which incorrectly detected eight *Culicoides* specimens, among them three members of the Obsoletus Group (*C. montanus*, *C. sanguisuga*, *C. sinanoensis*). Thus, incorrect detection within the subgenus *Avaritia* was significantly higher as compared to the newly developed mPCR, whereas cross-reactivity of the reference PCR primers to the members of the *Culicoides* subgenus *Culicoides* was comparatively low. However, the *C. scoticus* primer of the reference PCR (sco-COI-fwd) seems to be more specific than that of the new PCR, as it did not detect *C. scoticus* clade 2-DNA and showed less cross-reactivity to the other *Culicoides* species and haplotypes (n=2).

**Table 3.**
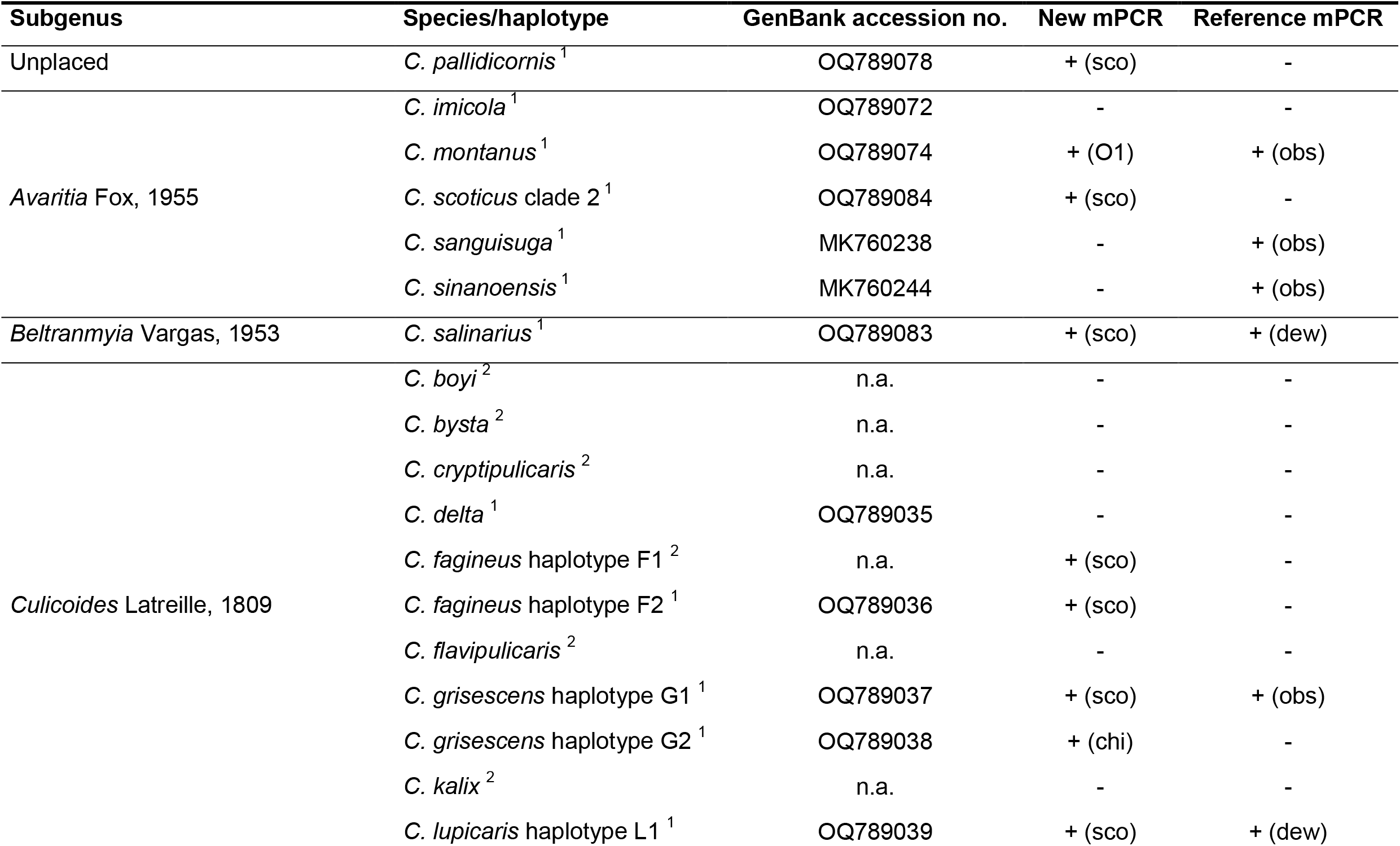
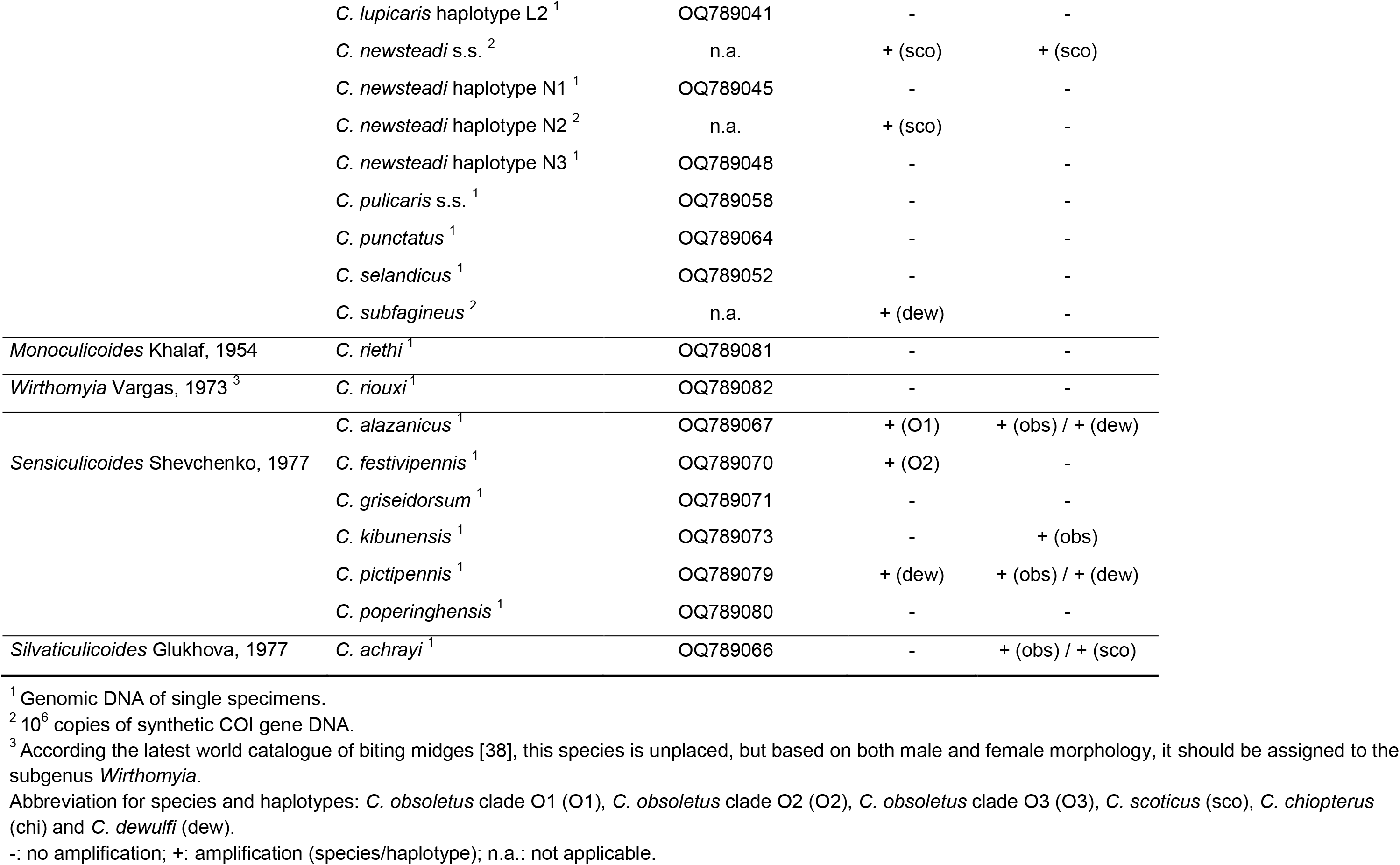
Cross-reactivity of the newly developed mPCR against 36 *Culicoides* species and haplotypes belonging to eight *Culicoides* subgenera as compared to the reference PCR test [70]. One specimen per species or haplotype was tested.

In addition to the specificity testing within the genus *Culicoides*, both mPCRs were checked regarding their cross-reactivity to non-ceratopogonid Diptera collected with BG-Sentinel traps to evaluate if the developed PCR can be applied on unsorted field collections (**Table 4**). Similar to the non-target biting midge species, both PCR tests unspecifically detected several non-ceratopogonid specimens. The newly developed mPCR incorrectly detected seven out of fourteen insect species, with the *C. dewulfi*-specific primer (dew-COI-470F) exhibiting the lowest specificity (n=4). In the case of the reference PCR, the primers obs-COI-fwd (n=6) and dew-COI-fwd (n=4) revealed deficiencies in their specificity, and in total, nine out of 14 species of by-catch were erroneously detected. In three cases (*Nemotelus notatus*, *Sepsis violacea* and *Sphaerocera curvipes*), the PCR even showed unspecific amplification with more than one primer, and the analysis of *Camptocladius stercorarius* produced an amplicon longer than 500 base pairs, not matching with the fragment lengths specifically produced.

**Table 4.** Cross-reactivity of the newly developed mPCR against 14 non-ceratopogonid dipteran species (possible by-catch in UV-light traps) as compared to the reference PCR test.

| Genus/species | GenBank accession no. | New mPCR | Reference mPCR |
| --- | --- | --- | --- |
| *Alluaudomyia* spec. | PP110213 | + (sco) | + (obs) |
| *Camptocladius stercorarius* | PP110214 | + (O3) | + (> 500 bp) |
| *Chironomus lugubris* | PP110215 | + (dew) | + (dew) |
| *Clogmia albipunctata* | PP110216 | - | + (obs) |
| *Forcipomyia* spec. | PP110217 | - | - |
| *Nemotelus notatus* | PP110218 | + (O1) / + (dew) | + (obs) / + (dew) |
| *Nilotanypus dubius* | PP110219 | - | - |
| *Physiphora alceae* | PP110220 | - | + (obs) |
| *Psychoda cinerea* | PP110221 | - | - |
| *Sepsis violacea* | PP110222 | + (dew) | + (sco) / + (dew) |
| *Smittia* spec. | PP110223 | + (dew) | + (dew) |
| *Spelobia luteilabris* | PP110224 | - | + (obs) |
| *Sphaerocera curvipes* | PP110225 | + (sco) | + (obs) / + (chi) |
| *Tephrochlamys rufiventris* | PP110226 | - | - |

## Discussion

In Europe, biting midge species of the wide-spread ‘Obsoletus/Scoticus Complex’, as well as *C. chiopterus* and *C. dewulfi*, appear to be key players in the transmission of the causative agents of epizootic BT and Schmallenberg diseases [75]. In the past years, three genetic variants of *C. obsoletus* and two clades of *C. scoticus* have been described, of which no unique biological details are known. Based on their relationship to other putative vector taxa, it is conceivable that they be competent virus vectors as well, having distinct ecological requirements, which demand selective management approaches. Such important biological issues could not be addressed so far due to the lack of identification assays.

In previous studies, the COI gene has proven to be a suitable marker for the differentiation of species of important insect taxa, including biting midges of the *Culicoides* subgenera *Culicoides* [41,61,54] and *Avaritia* [41,55–57], whereas in some biting midge subgenera (e.g. *Beltranmyia*, *Monoculicoides* and *Sensiculicoides*), the COI gene variances appear to be not adequate for species determination [41,76]. Generally, the utilization of mitochondrial genes for species delimitation is controversely discussed [e.g. 77-90]. Indeed, mitochondria are present in large copy numbers, and their genome is usually composed of conserved and variable regions [58–60], enabling molecular discrimination. Due to maternal inheritance, however, their genetic diversity is exclusively driven by the females [91] and not at all affected by the males. Interestingly, mitochondrial gene transfer is influenced by factors outside the individual’s evolutionary group, such as microbiota composition [91], and possible across genera or even biological kingdoms [77,92]. Certainly, the mentioned limitations are not to be dismissed, but could be remedied by the utilization of nuclear-coded gene loci [91] or the use of more than one genetic marker, as demonstrated by Mignotte et al. [36] who applied a multi-marker approach to revise the taxonomy of the Obsoletus/Scoticus Complex. This multi-marker strategy for an advanced PCR development could not be pursued in this study, due to the lack of sequence data of the relevant genes for the *Culicoides* species of interest. A sufficient number of GenBank entries was only available for the COI gene which, in consequence, was successfully used to develop a PCR test for species of the subgenus *Avaritia*, including recently described genetic variants, similarly to the scientific work by Dähn et al. [61] on the *Culicoides* subgenus *Culicoides*. The results of the present study confirm the suitability of the COI marker for species discrimination within the subgenus *Avaritia* [41,55–57] and underline the importance of open access genome data bases. However, initial sequence analysis turned out to be very challenging and demonstrated the limitations of such data repositories. The use of synonyms for one and the same culicoid species [38] and the numerous aliases of the mitochondrial cytochrome c oxidase subunit I gene (https://www.genecards.org/cgi-bin/carddisp.pl?gene=MT-CO1, last accessed on 16 August 2023) required extensive work to find suitable sequence regions. Once they were found, the direct comparison of species-specific sequences revealed incongruent results, suggesting a significant number of incorrect entries (e.g. up to 19.7% for *C. scoticus* clade 1). Such deficiencies in GenBank, which probably result from morphological misidentifications, synonymizations or the disability of the COI gene to distinguish respective species [93], have already been noted by other authors [41,61,94,95]. Hence, a more rigorous naming convention and cross-checking of submitted sequences as well as regular updating of GenBank entries through qualified experts is recommended. Considering that classical entomologists are at risk of becoming an ‘extinct species’ [61] and the considerable efforts necessary to maintain the continuously growing number of submitted sequences, the implementation of artificial intelligence (AI) could help manage these tasks.

Although the identification and re-evaluation of dubious COI sequences had to be performed without such tools in this study, the applied approach significantly facilitated the detection of such sequences and frequently resulted in accurate identifications, unveiling new perspectives on the taxonomy and abundance of the biting midge species. The interpretation of culicoid sequence data without additional morphological expertise, however, turned out to be challenging. This complexity arises from the natural presence of genetic variations (haplotypes) within each species, which further complicated the already problematic task of recognizing species borders. However, a procedure was found to generate consensus sequences with high intraspecific similarity, and their subsequent comparison revealed both new insights in the taxonomy of the subgenus *Avaritia* and genetic differences between taxa which could be used to design taxon-specific primers. For instance, the data analysis revealed only 2.8% sequence divergence between *C. obsoletus* clade O1 and *C. montanus*, and 2.9% between the two clades of *C. scoticus*, which is much less than the interspecific divergence determined for the other species and haplotypes of the subgenus (7.9%-23.7%). Although the delineation of species is inherently challenging [78,96–98], the results strongly indicate that, from a genetic point of view, these entities represent genetic variations rather than distinct species, maybe due to recent speciation events, supporting suggestions in a previous study comparing COI data of specimens morphologically identified as *C. obsoletus* and *C. montanus* [36]. According to the above mentioned doubts regarding the use of mitochondrial genes and the proposed alternative use of nuclear gene-loci for taxonomic investigations, the GenBank was browsed for nuclear ribosomal ITS1/ITS2 (rDNA) sequences of specimens, whose COI sequence were used in the framework of this study and could be assigned either to *C. obsoletus* clade O1 or *C. montanus*. A comparison of these sequences were meant to reveal if our conclusions regarding the taxonomic status of *C. montanus* were supportable. A comparison of the few rDNA sequences of *C. montanus* (n=3, MK893026-28) and *C. obsoletus* (n=6, MK893032-37) specimens found [44] also showed very low interspecific divergence (1.1%). Similar results (0.8% divergence between *C. obsoletus* and *C. montanus*), apparently supporting our hypothesis, were achieved through analysis of sequence data of the 16S rDNA gene in another study [36].

In contrast to the close relationship of *C. obsoletus* clade O1 and *C. montanus*, the genetic divergence between the three clades of *C. obsoletus* was comparatively high (> 9.3%), arguing for species status. This demand is supported by the observation that *C. obsoletus* clade O1 showed lower genetic distance to *C. sanguisuga* and *C. sinanoensis*, two accepted species of the Obsoletus Group, than to the *C. obsoletus* clades O2 and O3. As already discussed in previous studies [36,41], the high COI sequence similarity of *C. obsoletus* s.s. and *C. obsoletus* clade O1 obtained in the present study suggest synonymity. By contrast, the assumption that *C. obsoletus* clade O3 should be regarded synonymous to *C. gornostaevae* [36] could neither be confirmed nor rejected, as no study has yet analysed individuals morphologically and genetically at the same time. In the case of *C. obsoletus* clade O2, two possible scenarios are imaginable. Firstly, it represents one of the described valid species of the subgenus *Avaritia* or, secondly, it represents a previously unknown parametric species. As for *C. dewulfi*, the results of COI gene analysis seem to confirm its position outside the Obsoletus Group [35,40–44], whereas the status of *C. chiopterus* remains ambiguous [35,43] with regard to the calculated interspecific divergences from the other Obsoletus Group members, which were not significantly higher than the divergence from *C. alachua*. Nevertheless, it should be noted that genetic analysis alone is not sufficient to define species boundaries or perform taxonomic revisions, which are essential for the development of molecular tests. Hence, the combination of morphological, molecular, and ecological data is needed to solve such complex issues [99], requiring classical taxonomists as well as experienced molecular biologists.

Due to the similarity of the COI sequence of *C. obsoletus* clade O1 and *C. montanus*, no specific forward primers could be designed in this study that would not detect the other taxon. Hence, samples producing a PCR amplicon of 120 bp with the presented PCR are to be designated ‘*C. obsoletus* clade O1/*C. montanus*’ and have to be further analysed by sequencing. The same applies to the *C. scoticus*-primer, which produces a 317 bp-PCR fragment with both the DNAs of *C. scoticus* clade 1 and clade 2. Interestingly, despite the low genetic distance, PCR tests based on the ribosomal ITS1 [65] and ITS2 [68] regions have been developed for *C. obsoletus* and *C. montanus*, which could be used alternatively, but the reliability of these tests has not been checked in this study. By contrast, primer development was successful in the present study for the COI differentiation of *C. chiopterus*, *C. dewulfi* and the three clades of *C. obsoletus*.

Based on the findings obtained, it is supposed that *C. montanus* constitutes a genetic variant of *C. obsoletus* clade O1 and that *C. scoticus* clade 2 represents a haplotype of *C. scoticus* clade 1 (thereby making the distinction of *C. scoticus* clades obsolete). Considering this, the designed primers exhibit the ability of accurately identifying their respective target taxa within the subgenus *Avaritia*, as could be shown through the analysis of field-caught biting midges. This precise detection is achieved with a single-tube multiplex PCR, which was designed to be carried out in parallel (i.e. with the same cycling conditions) with the recently described multiplex PCRs for the identification of the presently known members of the *Culicoides* subgenus *Culicoides* [61].

Due to observed cross-reactivity of some of the newly designed forward primers with *Culicoides* species not belonging to the subgenus *Avaritia*, morphological pre-identification of the biting midges to group level is mandatory. Simply because of the huge number of taxa that had to be considered during PCR development, absolute primer specificity is extremely difficult to achieve, based on available sequence data. Especially in the case of the *C. scoticus*-primer (sco-COI-317F), genetic variations in the primer annealing region are incapable of preventing primer binding to genomic DNA of nine other *Culicoides* species belonging to different subgenera and two non-ceratopogonid dipteran species. A comparison of the primer binding site sequences of respective species with those of *C. scoticus* revealed nucleotide differences in one to four positions (c.f. **Supplementary Table S4**), primarily in the middle or at the 5’-end. Such mismatches did not lead to the complete loss of primer binding capacity, although a high annealing temperature and a hot-start Taq polymerase were used in this study to reduce unspecific primer annealing [72,73]. As suggested by Dähn et al. [61], the targeted insertion of mismatch bases was also considered during PCR development and actually increased the specificity of the primer dew-COI-470F within the subgenus *Avaritia* significantly, but did not prevent unspecific detection of other *Culicoides* and non-ceratopogonid insects. In some cases, more than four mismatches in the primer binding region were insufficient to avoid unspecific binding (**Supplementary Table S4**) although the opposite should be expected. Similar observations were made and discussed during the development of multiplex PCRs for the subgenus *Culicoides* [61].

Despite the observed cross-reactivity of some PCR primers, morphologically pre-identified species of the subgenus *Avaritia* were identified with high accuracy. Advantages of the developed PCR for routine identification of subgenus *Avaritia* biting midge species over the commonly used COI barcoding approach are that it (i) does not detect the donor of a recently taken blood-meal of engorged females (due to the utilisation of culicoid-specific universal forward primers), and (ii) is capable of evaluating the presence or absence of adult specimens of the West Palaearctic *Culicoides* subgenus *Avaritia* members (including recently discovered clades of *C. obsoletus*) in a pool of up to 50 specimens. With respect to the assumption of Meiswinkel et al. [37], that *C. obsoletus* ’clade dark’ might be *C. gornostaevae*, the developed PCR approach could be the first molecular test that specifically detects this putative vector species and help answer open questions regarding habitat preference and other ecological traits, which is a prerequisite for an effective vector management.

## Conclusions

The multiplex PCR approach developed in this study is capable of detecting all putative vector species of the subgenus *Avaritia* belonging to the West Palaearctic biting midge fauna, and for the first time enables the differentiation of recently discovered clades of *C. obsoletus*, whose role in virus transmission is unknown. Although the PCR was mainly validated with biting midge specimens collected in Germany, primer design was achieved through bioinformatic analysis of all COI sequences available from GenBank, which highlights the importance of such databases. The derived specific consensus sequences showed high intraspecific similarities suggesting that the test might be applicable to a much broader geographic area. The results presented in this study once again revealed the potential of the COI marker for the development of molecular tools and resulted in an easy-to-use multiplex PCR becoming now available to get a deeper insight into the distribution of the different West Palaearctic members of the subgenus *Avaritia* and possibly contributing to improve future risk assessment of culicoid-borne diseases, such as bluetongue, Schmallenberg disease, African horse sickness or epizootic hemorrhagic disease of cervids. The causative agent of the latter has only recently invaded Europe [100] and has been shown to be able to disseminate in both *C. obsoletus* and *C. scoticus*, suggesting vector competence, although genotyping of the biting midge species was not performed [101]. By contrast, the newly developed PCR was successfully used in practice for identifying biting midge vector species involved in the recent BTV-3 outbreak in western Europe [102].

## Acknowledgments

We thank our colleagues Sarah Groschupp, Christin Henneberg, Maria Will, Nathalie Richter and Sebastian Schuran from the Leibniz Centre for Agricultural Landscape Research (ZALF) for the comprehensive work during the collection and morphological pre-identification of biting midges used in this study. We are also grateful to Maxi Uecker and Freja Pfirschke (FLI) for the excellent technical assistance regarding the molecular biological work.

## Data Availability Statement

New data created in this study are deposited in GenBank (https:// www.ncbi.nlm.nih.gov/genbank/) with the following accession numbers: *C. obsoletus* clade O1: OQ941500-OQ941537; *C. obsoletus* clade O2: OQ941538-OQ941560 and PP110209-PP110212; *C. obsoletus* clade O3: OQ941561-OQ941562; *C. scoticus* clade 1: OQ941563-OQ941572; *C. chiopterus*: OQ941573; *C. dewulfi*: OQ941574-OQ941576; *Alluaudomyia* spec.: PP110213; *Camptocladius stercorarius*: PP110214; *Chironomus lugubris*: PP110215; *Clogmia albipunctata*: PP110216; *Forcipomyia* spec.: PP110217; *Nemotelus notatus*: PP110218; *Nilotanypus dubius*: PP110219; *Physiphora alceae*: PP110220; *Psychoda cinerea*: PP110221; *Sepsis violacea*: PP110222; *Smittia* spec.: PP110223; *Spelobia luteilabris*: PP110224; *Sphaerocera curvipes*: PP110225; *Tephrochlamys rufiventris*: PP110226.

## Supplementary Material

**Supplementary Table S1:** Analyzed GenBank entries and produced consensus sequences used for the development of forward primers specific for the different taxa of the subgenus *Avaritia*.

**Supplementary Table S2:** Sequences of synthetic COI genes of subgenus *Culicoides* and subgenus *Avaritia* taxa used in this study.

**Supplementary Table S3:** List of all forward primers tested in this study.

**Supplementary Table S4:** Cross-talk of newly designed primers with other *Culicoides* spec. and non-ceratopogonid insect species.

## Notes

### Competing Interest Statement

The authors have declared no competing interest.

